# A 3D-printed pump-free multi-organ-on-a-chip platform for modeling the intestine–liver–muscle axis

**DOI:** 10.1101/2025.09.02.673650

**Authors:** Rodi Kado Abdalkader, Takuya Fujita

**Author notes:** Earlier known as Rodi Abdalkader.

## Abstract

The intestine–liver–muscle axis plays an essential role in drugs and nutrients absorption, metabolism, and energy balance. Yet in vitro models capable of recapitulating this inter-organ communication remain limited. Here, we present a pump-free, 3D-printed multi-organ-on-a-chip device that enables dynamic co-culture of Caco-2 intestinal epithelial cells, HepG2 hepatocytes, and primary human skeletal myoblasts (HSkM) under gravity-driven oscillatory flow. The device consists of five interconnected chambers designed to accommodate Transwell cell culture inserts for intestine and muscle compartments and hydrogel-embedded hepatocyte spheroids in the central hepatic compartment. The device was fabricated by low-cost fused deposition modeling (FDM) using acrylonitrile butadiene styrene (ABS) polymers. Under dynamic rocking, oscillatory perfusion promoted inter-organ communication without the need for external pumps or complex tubing. Functional assessments revealed that dynamic co-culture significantly enhanced the functions of skeletal muscle, as indicated by increased myosin heavy chain expression and elevated lactate production, while HepG2 spheroids exhibited improved hepatic function with higher albumin expression compared with monoculture. Additionally, Caco-2 cells maintained stable tight junctions and transepithelial electrical resistance, demonstrating preserved intestinal barrier integrity under dynamic flow. These results establish the device as a versatile, accessible 3D-printed platform for modeling the intestine–liver–muscle axis and investigating metabolic cross-talk in drug discovery and disease modeling.

## Introduction

The human body maintains metabolic homeostasis through tightly coordinated interactions among multiple organ systems. Among these, the intestine, liver, and skeletal muscle form a central axis that governs nutrient absorption, energy metabolism, and the pharmacokinetics of orally administered drugs^1–3^. The intestine acts as the initial barrier for nutrients and xenobiotics, controlling absorption through its polarized epithelial lining and selectively regulating transport into the portal circulation ^4^. Following absorption, the liver serves as the primary site for first-pass metabolism, performing a wide range of enzymatic transformations that determine drug bioavailability and systemic exposure ^5,6^. Skeletal muscle, as the largest metabolic organ, is a major site of glucose and fatty acid utilization, responding to circulating metabolites and hormonal cues to maintain systemic energy balance ^7^. Disruption within this axis underlies a spectrum of clinical conditions, including insulin resistance, sarcopenia, and non-alcoholic fatty liver disease, highlighting the importance of accurately modeling inter-organ communication for both basic research and drug development^8,9^.

Conventional in vitro models, such as static monocultures or simple co-cultures, are limited in their ability to replicate the dynamic exchange of nutrients, hormones, and metabolites between organs ^10^. Transwell cell culture systems, where cells are cultured on semi-permeable membranes, provide a partial solution by mimicking tissue barriers and allowing selective transport, and they are widely used for evaluating intestinal permeability and epithelial function ^11^. However, traditional Transwell-based approaches remain static and fail to reproduce the dynamic fluidic conditions and multi-organ interactions that occur in vivo ^12^. Meanwhile, in vivo studies in animal models offer insights into systemic metabolism but are constrained by interspecies differences in transporter expression, enzyme activity, and tissue response, which often limit the predictive value of preclinical data for human outcomes ^13,14^.

Organ-on-a-chip (OOC) and microphysiological systems (MPS) have emerged as transformative platforms to bridge this gap, enabling the recapitulation of human tissue function under controlled microfluidic environments^15,16^. By integrating multiple tissue types into interconnected compartments, organ-on-a-chip systems allow the study of organ-to-organ communication, drug absorption, and metabolism with higher physiological relevance [19]. However, many existing OOC platforms rely on external peristaltic pumps or complex microfluidic controllers, which increase operational complexity, cost, and the technical barrier for routine use in standard laboratories ^17^. To overcome these limitations, we used our previously developed platforms ^18,19^as the basis for a 3D-printed, pump-free multi-organ-on-a-chip device that integrates Transwell-based culture insert compartments for intestine, liver, and skeletal muscle. The use of 3D printing enables rapid prototyping, low-cost production, and customizable design, making the platform accessible for widespread implementation. Moreover, the gravity-driven bi-directional flow generated by simple rocking eliminates the need for external pumps while maintaining physiologically relevant inter-organ exchange.

In this study, we present a gravity-driven, 3D-printed multi-organ-on-a-chip platform that models the intestine–liver–muscle axis. Caco-2 cells on Transwells cell culture inserts formed an intestinal barrier, human skeletal myoblasts provided a metabolically active muscle compartment, and HepG2 spheroids embedded in collagen/Matrigel served as the liver component for first-pass metabolism. The device, fabricated by low-cost 3D printing using acrylonitrile butadiene styrene (ABS) polymer with PDMS sealing, operates under gentle rocking to generate pump-free, bi-directional flow. By integrating Transwell cell culture inserts with dynamic microfluidics, the system offers a scalable and accessible platform for physiologically relevant co-culture for the study of inter-organ communication.

## Results

### Design and fabrication of a multi-organ-on-a-chip device

We first developed a 3D-printed multi-organ-on-a-chip device to enable gravity-driven co-culture of intestine, liver, and skeletal muscle tissues without the use of external pumps (**Figure 1a**). The device comprised five aligned circular chambers: two reservoirs at the ends, one central chamber for HepG2 hepatocytes, and two side chambers for Caco-2 intestinal and human skeletal myoblast (HSkM) cultures, all interconnected by a lower circulation channel. Arrows in the schematic indicate the bidirectional flow path generated by gentle rocking of the device at ±6°, which drives fluid oscillation and nutrient exchange between compartments.

**Figure 1.**
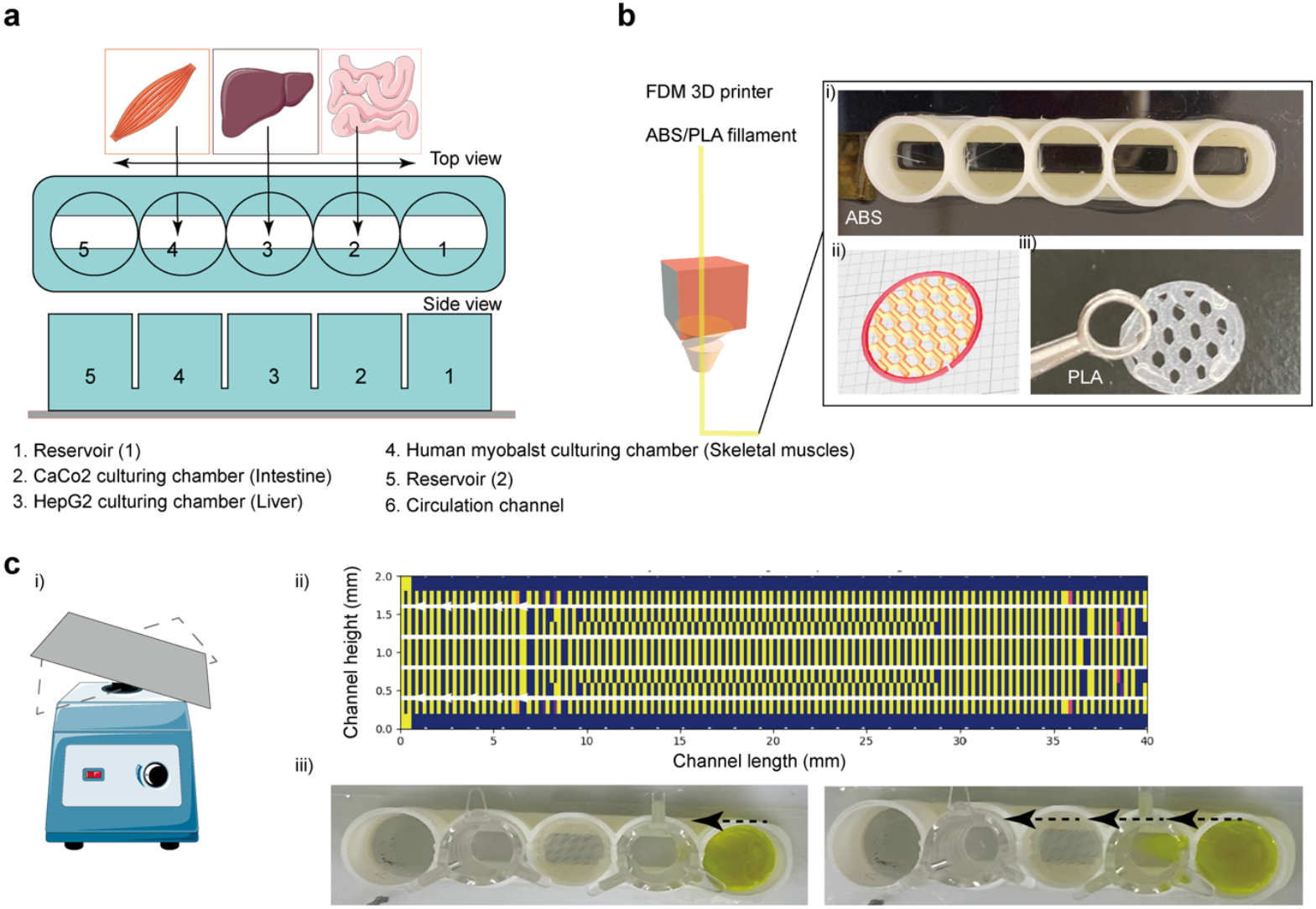
Design and fabrication of a 3D printed multi-organ-on-a-chip device for gravity driven co-culture of intestine, liver, and skeletal muscle tissues. **a**, Schematic of the device showing the arrangement of organ specific chambers and reservoirs: (1) inlet reservoir, (2) Caco-2 intestinal chamber, (3) HepG2 hepatic chamber, (4) human skeletal myoblast chamber, (5) outlet reservoir, and (6) connecting circulation channel. Arrows indicate the direction of gravity driven bidirectional flow during rocking. **b**, Fabrication of the device using fused deposition modeling (FDM) 3D printing. (i) Photograph of the printed ABS body with five aligned organ chambers. (ii) CAD design illustrating the internal lattice used to support hydrogel or ECM seeding. (iii) PLA lattice insert for hydrogel embedding and enhanced cell attachment. c, Dynamic perfusion setup and demonstration of oscillatory flow. (i) Device mounted on a rocking platform (±6°) to generate laminar bidirectional flow without external pumps. (ii) Flow profile visualization along the connecting channel, showing 2 D projection of dye distribution over the channel height and length. (iii) Experimental observation of tracer (fluorescein sodium) oscillating through the interconnected chambers, confirming passive mixing and nutrient transport across the multi-organ system.

The device body was fabricated using FDM 3D printing with ABS for the main structure and PLA inserts for hydrogel support (**Figure 1b**). The modular design included a lattice structure to secure collagen/Matrigel droplets for hepatocyte spheroid culture, and flat surfaces compatible with Transwell cell culture inserts for the intestinal and muscle compartments. Experimental setup on a rocking platform confirmed the generation of oscillatory fluid movement, which could be visualized using tracer dye (**Figure 1c**) (**Supplementary figure 1**) **(Supplementary video 1)**. Back-and-forth dye oscillation demonstrated that gravity-driven flow is sufficient to perfuse all chambers, enabling multi-organ communication without the need for complex microfluidic pumps.

### Cell preparation and integration into the device

To establish a physiologically relevant multi-organ system, we optimized the timing and configuration of cell preparation prior to integration (**Figure 2a**). Caco-2 cells were pre cultured in Transwell cell culture inserts for 14 days to form polarized monolayers. HepG2 were encapsulated in collagen/Matrigel to generate 3D spheroids over 5 days, providing a liver like 3D microenvironment. HSkM cells were pre cultured for 2 days to establish a confluent monolayer in Transwells before assembly into the chip. At day 0, all three tissues were transferred to the device, where they were co cultured under gravity driven oscillatory flow for 3 days to enable inter organ communication. Representative images show the morphology and maturity of each tissue prior to device integration (**Figure 2b**). HSkM monolayers exhibited elongated, confluent morphology after 2 days, while HepG2 cells formed compact 3D spheroids within hydrogel droplets by day 5. Caco-2 monolayers displayed a uniform epithelial layer consistent with functional barrier formation after 14 days. The schematic below the images illustrates how these preconditioned tissues were spatially arranged in the device, with intestinal and muscle compartments in Transwell cell culture inserts and the hepatic compartment embedded in the central hydrogel chamber. This preparation strategy allowed all three tissues to be introduced in a functionally ready state, supporting immediate cross talk upon initiation of dynamic culture.

**Figure 2.**
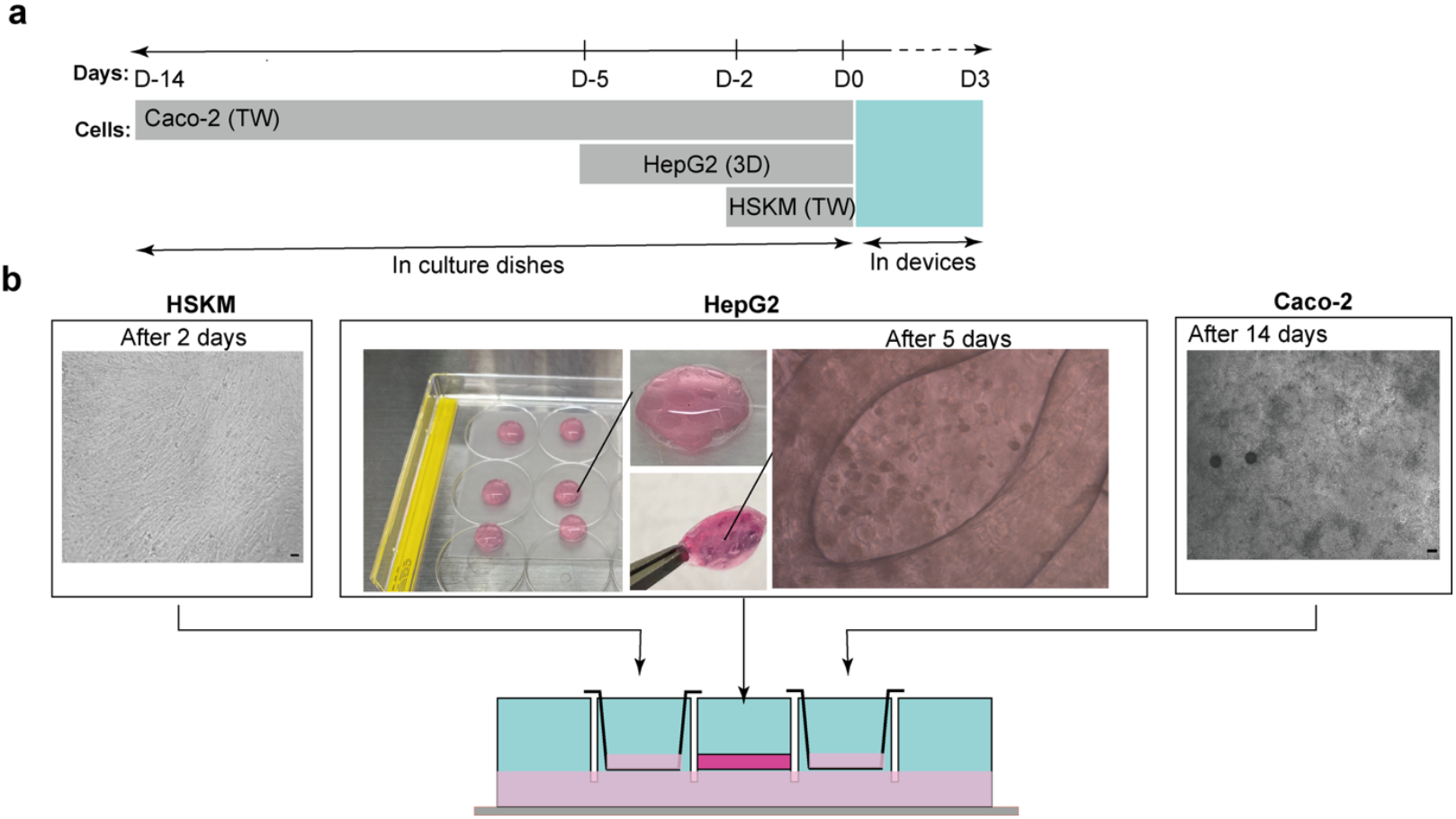
Cell preparation and integration workflow for multi-organ co-culture in the 3D-printed device. **a**, Timeline of cell preparation and device assembly. Caco-2 cells were pre-cultured in Transwell cell culture inserts for 14 days to form polarized intestinal monolayers, while HepG2 hepatocytes were aggregated into 3D spheroids over 5 days in collagen/Matrigel droplets, and primary human skeletal myoblasts (HSkM) were pre-cultured for 2 days in Transwells. At day 0 (D0), the three tissue types were integrated into the device for 3 days of dynamic co-culture under gravity-driven bidirectional flow. **b**, Representative images of individual tissue compartments prior to integration: **HSkM (left):** Confluent myoblast monolayer after 2 days of culture. **HepG2 (center):** Collagen/Matrigel spheroids after 5 days of culture, shown as whole droplets (inset) and under brightfield microscopy. **Caco-2 (right):** Polarized monolayer in Transwell cell culture insert after 14 days. The schematic below illustrates the spatial arrangement of the three tissues in the multi-organ-on-a-chip device, with Transwell cell culture inserts for intestinal and muscle compartments and a central hydrogel chamber for hepatic spheroids.

### Dynamic co-culture enhances skeletal muscle function and metabolic activity

To evaluate skeletal muscle function, HSkM cells were cultured as monocultures (Mono), or co-cultured in the device under static (CO_ST) or dynamic (CO_DY) conditions (**Figure 3a**). Immunofluorescence staining for myosin heavy chain (MYH) revealed more extensive myotube formation in co-culture compared to monoculture, with the strongest expression observed under dynamic conditions. Quantitative analysis of MYH expression confirmed a significant increase in CO_DY compared with Mono and CO_ST cultures (**Figure 3b**). In parallel, lactate levels measured in the circulation channel were significantly higher in CO_DY compared to CO_ST, indicating enhanced glycolytic activity under dynamic flow (**Figure 3c**). Together, these results demonstrate that dynamic co-culture promotes both differentiation and metabolic activity of HSkM.

**Figure 3.**
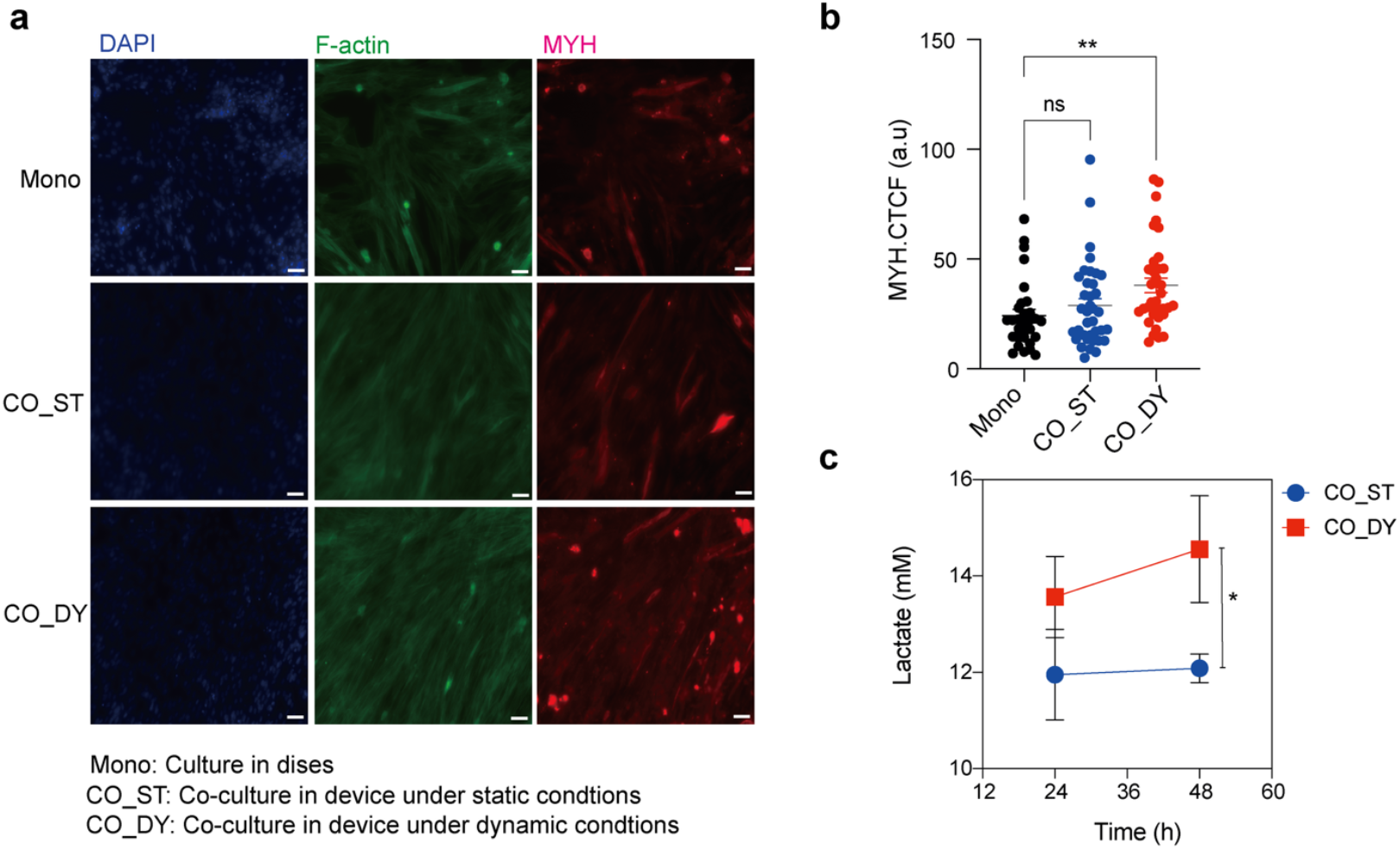
Skeletal muscle differentiation and metabolic activity under mono- and co-culture conditions. **a**, Immunofluorescence staining of human skeletal myoblasts cultured in monoculture (Mono), in the multi-organ device under static conditions (CO_ST), and in the device under dynamic rocking conditions (CO_DY) at day 3. Nuclei (DAPI, blue), F-actin (green), and myosin heavy chain (MYH, red) are shown. Scale bars, 50 µm. **b**, Quantification of MYH expression by corrected total cell fluorescence (CTCF) in Mono, CO_ST, and CO_DY cultures. Dynamic co-culture significantly enhanced MYH expression compared with static and monoculture conditions (one-way ANOVA with Dunnett’s post hoc test, **p < 0.01; ns = not significant). **c**, Lactate concentration measured in the circulation channel under static (ST) and dynamic (DY) co-culture conditions at 24–48 h. Dynamic culture resulted in significantly higher lactate release compared with static conditions (unpaired Student’s t-test, *p < 0.05). Data are presented as mean ± SEM.

### Hepatic function of HepG2 spheroids is improved under dynamic co-culture

HepG2 cells encapsulated in collagen/Matrigel scaffolds maintained compact spheroid morphology across all conditions (**Figure 4a**). However, immunofluorescence staining revealed stronger albumin (ALB) expression in co-cultures compared with monocultures, with the highest expression observed in CO_DY (**Figure 4b**). Quantification of ALB intensity confirmed that both static and dynamic co-culture significantly enhanced hepatic function relative to monoculture, with dynamic flow further increasing albumin expression (**Figure 4c**). These findings indicate that co-culture with other tissues in the device, particularly under dynamic perfusion, supports improved hepatic activity of HepG2 spheroids.

**Figure 4.**
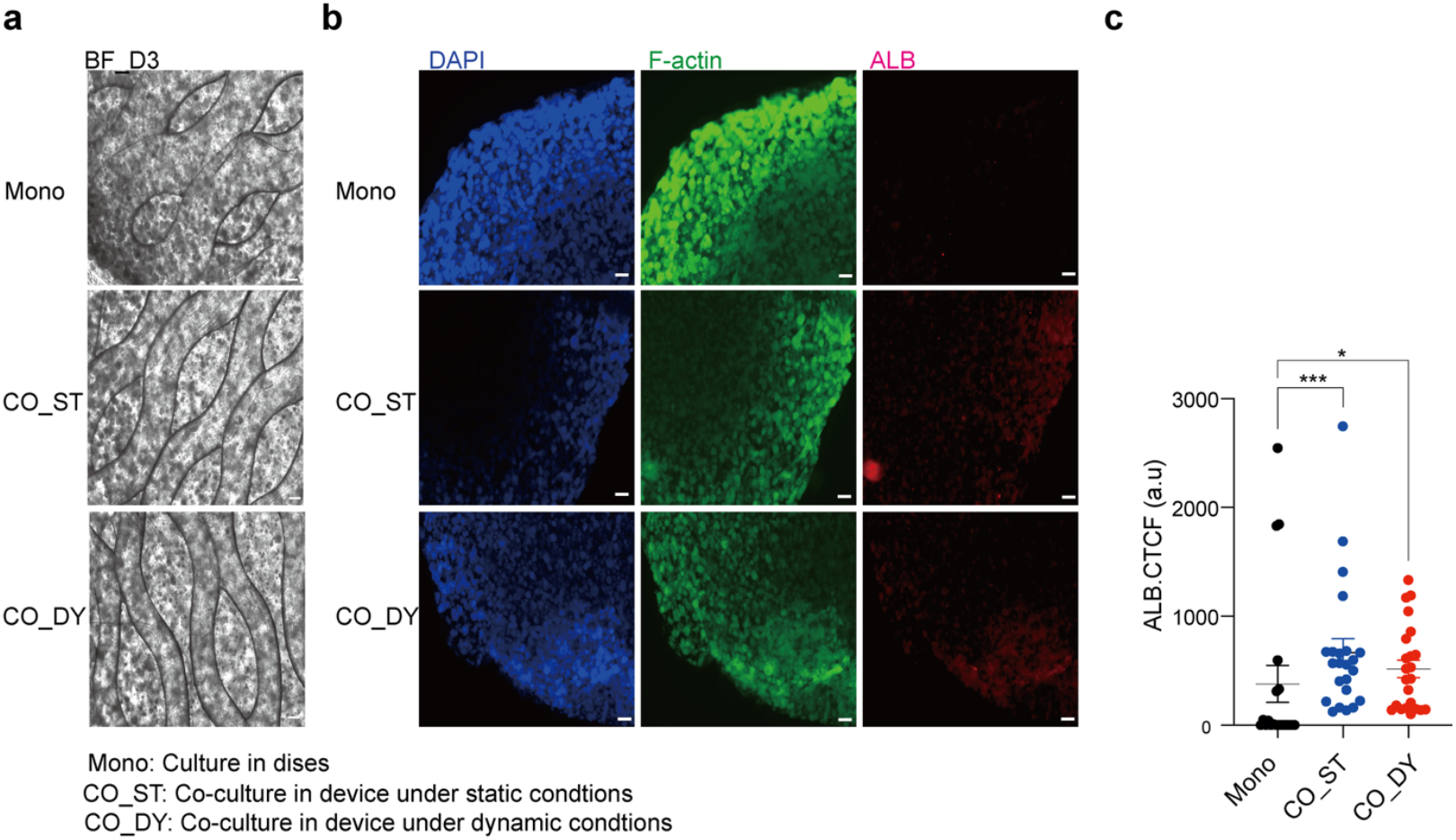
Morphology and functional activity of HepG2 cells under mono- and co-culture conditions. **a**, Brightfield images of HepG2 spheroids seeded on PLA scaffolds after 3 days of culture in monoculture (Mono), in the multi-organ device under static conditions (CO_ST), and in the device under dynamic rocking conditions (CO_DY). Scale bars, 200 µm. **b**, Immunofluorescence staining of HepG2 spheroids showing nuclei (DAPI, blue), F-actin (green), and albumin (ALB, red). Albumin expression was markedly enhanced under co-culture conditions, with the strongest expression observed under dynamic flow. Scale bars, 200 µm. **c**, Quantification of ALB expression by corrected total cell fluorescence (CTCF) in Mono, CO_ST, and CO_DY conditions. Both static and dynamic co-cultures significantly increased ALB expression compared with monoculture (one-way ANOVA with Dunnett’s post hoc test; ***p < 0.001, *p < 0.05). Data are presented as mean ± SEM.

### Caco-2 intestinal barrier integrity is preserved under dynamic conditions

To examine whether dynamic rocking influenced intestinal barrier function, Caco-2 monolayers were analyzed by ZO-1 immunofluorescence and transepithelial electrical resistance (TEER). ZO-1 staining revealed continuous tight junctions under both CO_ST and CO_DY conditions (**Figure 5a**). Quantification of ZO-1expression showed no significant differences between static and dynamic co-culture (**Figure 5b**). TEER values were consistently above 200 Ω·cm^2^ in both conditions, with no significant changes between CO_ST and CO_DY (**Figure 5c**). These results indicate that dynamic rocking preserved the epithelial barrier properties of Caco-2 monolayers.

**Figure 5.**
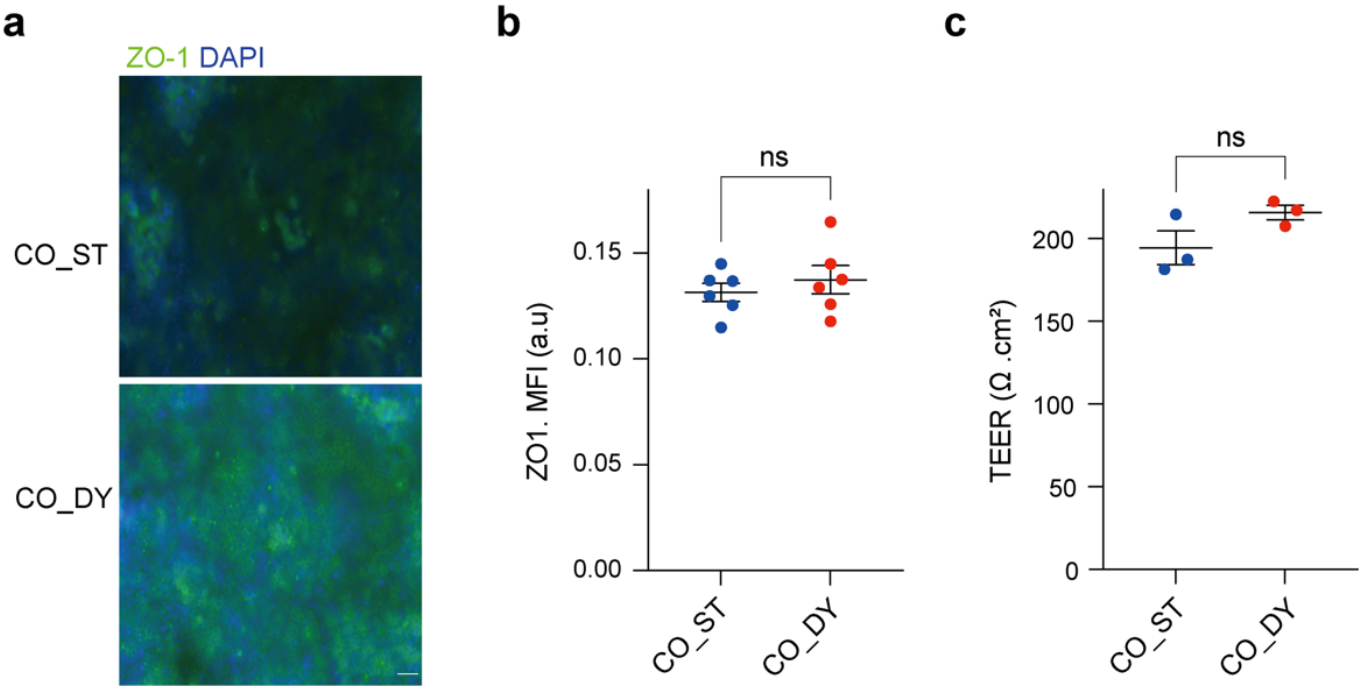
Barrier integrity of Caco-2 monolayers under static and dynamic co-culture conditions. **a**, Immunofluorescence staining of Caco-2 cells for tight junction; ZO-1 (green) and nuclei (DAPI, blue) in the multi-organ device under static (CO_ST) and dynamic rocking (CO_DY) conditions. Scale bars, 50 µm. **b**, Quantification of ZO-1 expression by mean fluorescence intensity (MFI) showing no significant difference between static and dynamic conditions (ns). **c**, Transepithelial electrical resistance (TEER) measurements of Caco-2 monolayers under static (CO_ST) and dynamic (CO_DY) conditions. No significant differences were observed (ns). Data are presented as mean ± SEM.

## Discussion

In this study, we developed and evaluated a 3D-printed, pump-free multi-organ-on-a-chip device that integrates intestinal, hepatic, and skeletal muscle tissues under gravity-driven oscillatory flow. By combining Transwell cell culture inserts for barrier and muscle compartments with hydrogel-embedded hepatocyte spheroids, this system recapitulates key aspects of inter-organ communication while maintaining accessibility and low operational complexity.

Our results demonstrate that dynamic perfusion enhances muscle function and metabolic activity, supports improved hepatic function, and preserves intestinal barrier integrity. The observed increase in myosin heavy chain expression and lactate production under dynamic co-culture highlights the importance of mechanical cues and inter-organ communication in regulating skeletal muscle function. Traditional static culture often fails to reproduce the contractile phenotype or metabolic activity of skeletal muscle, limiting its translational relevance. Our results align with previous reports showing that microfluidic perfusion promotes nutrient exchange and waste clearance, thereby enhancing cell functions^20^.^21^ Importantly, by eliminating external pumps and relying on gravity-driven oscillation, our approach lowers the technical barrier for implementing dynamic culture.

Albumin secretion is a key indicator of hepatocyte function, and our findings demonstrate that both static and dynamic co-culture with intestinal and muscle compartments enhanced HepG2 performance compared with monoculture^22,23^. This improvement is consistent with the notion that hepatic cells depend on paracrine factors and metabolic inputs from other organs. The further enhancement under dynamic flow suggests that oscillatory perfusion facilitates efficient metabolite exchange, mimicking features of portal circulation. While HepG2 cells are an immortalized cell line and do not fully capture primary hepatocyte physiology, the increased albumin expression highlights the ability of this platform to promote hepatic functionality in a multi-organ context.

The preservation of tight junction protein ZO-1 and stable TEER values under dynamic culture indicate that intestinal epithelial integrity was not compromised by oscillatory flow. This is significant because barrier disruption often confounds co-culture systems, especially under mechanical stimulation. Our results suggest that the combination of Transwell cell culture inserts and gentle rocking provides a robust model for studying nutrient and xenobiotic absorption without compromising epithelial physiology.

Most existing multi-organ-on-a-chip platforms rely on complex microfluidic controllers or peristaltic pumps to achieve inter-organ communication. While powerful, such systems are often costly, technically demanding, and difficult to implement in non-specialist laboratories. In contrast, our 3D-printed device can be fabricated at low cost using widely available materials (ABS, PLA, TPU) and operated with a simple rocking platform. The modular design also enables rapid prototyping and adaptation to different organ configurations. These features address a major gap between cutting-edge organ-chip technologies and their practical adoption for routine use in drug testing, disease modeling, and academic research.

The ability to simultaneously culture intestine, liver, and skeletal muscle tissues under dynamic conditions opens new opportunities for investigating metabolic diseases such as insulin resistance, sarcopenia, and non-alcoholic fatty liver disease. Nonetheless, several limitations remain: we used immortalized cell lines (Caco-2, HepG2), which do not fully reproduce the complexity of primary or iPSC-derived tissues; the duration of co-culture was limited to short-term (3 days), and longer experiments will be necessary to evaluate stability, maturation, and chronic responses; and while we measured basic metabolic activity, a comprehensive assessment of drug metabolism, cytochrome P450 activity, and transport kinetics was beyond the scope of this study but will be essential for advancing toward pharmacokinetic and toxicological applications. Future work should explore the integration of primary human cells or iPSC-derived lineages to enhance physiological relevance, as well as the incorporation of biosensors for real-time monitoring of metabolites, cytokines, and contractile activity.

## Conclusion

In this study we demonstrate that a low-cost, 3D-printed multi-organ-on-a-chip device can successfully support intestine–liver–muscle co-culture under gravity-driven perfusion. Dynamic flow enhanced muscle differentiation and hepatic function without compromising intestinal barrier integrity, validating the system as a versatile and physiologically relevant platform. By bridging accessibility and functionality, this approach expands the potential of organ-on-a-chip technology for both basic research and translational applications in drug discovery and metabolic disease modeling.

## Methods

### Device fabrication and characterization

The multi-organ-on-a-chip device was fabricated following our previously reported method ^18,19^ using fused deposition modeling (FDM) 3D printing (Anycubic) with 1.75 mm acrylonitrile butadiene styrene (ABS) filaments. Computer-aided design (CAD) models were generated using Tinkercad (https://www.tinkercad.com) (**Supplementary Figure 1**), and STL files were processed using Ultimaker Cura software. Printing parameters were set to a layer height of 0.01–0.02 mm and a shell thickness of 2–10 layers. Printed devices were adhered to PET clear dishes using PDMS, while Polylactic acid (PLA) lattice inserts were printed separately to support hydrogel encapsulation. After printing, excess edges were trimmed, and devices were stored at room temperature until use. To assess fluid dynamics within the microchannels, fluorescein sodium (1 mg mL^-1^, Nacalai) was perfused under rocking conditions, and oscillatory flow was confirmed using bright-field microscopy. To support the experimental flow visualization, a two-dimensional simulation of oscillatory fluid motion in the circulation channel (40 × 4 × 2 mm; L × W × H) was performed. The flow was modeled as laminar with a parabolic velocity profile, and the mean velocity was set to oscillate sinusoidally with time to mimic platform rocking (±6°, 10 cycles min^-1^, peak velocity ≈ 5 mm s^-1^ as measured experimentally). Tracer transport was described by the advection–diffusion equation using the diffusion coefficient of fluorescein in water (4×10^−10^ m^2^ s^-1^)^24^. The equation was solved numerically with a finite-difference method in Python (NumPy, Matplotlib). No-flux boundary conditions were applied at the channel walls. Simulation output was rendered as animations showing tracer oscillation and mixing (**Supplementary Code S1**).

### Cell culture and multi-organ assembly

Caco-2 cells (RIKEN BRC) were maintained in DMEM (Wako) supplemented with 10% fetal bovine serum (FBS; Thermo Fisher Scientific), 1% non-essential amino acids (Nacalai), and 1% penicillin–streptomycin (Wako). Cells were seeded at 0.5 × 10^5^ cells per insert onto 24-well polyester Transwell inserts (0.4 µm pore size, ThinCert; Greiner Bio-One), with medium supplied apically (0.25 mL) and basolaterally (0.75 mL). Medium was exchanged every other day. After 14 days, Caco-2 cells formed polarized epithelial monolayers, confirmed by stable transepithelial electrical resistance (TEER ≥ 200 Ω·cm^2^).

Human primary normal skeletal myoblasts (HSkM; Gibco) were cultured in DMEM (Wako) supplemented with 2% horse serum (Thermo Fisher Scientific) to induce differentiation. Cells were seeded at a density of 2.4 × 10^4^ per Transwell insert and incubated for 24 h at 37 °C with 5% CO_2_. The culture medium was replaced daily with fresh differentiation medium. After 2 days, confluent monolayers were obtained and used for device integration.

HepG2 cells (RIKEN BRC) were suspended in a collagen I/Matrigel mixture (Cellmatrix I-A, Nitta Gelatin / Geltrex, Thermo Fisher Scientific) at a 9:1 ratio (final collagen concentration 3 mg mL^-1^) at a density of 4 × 10^5^ cells mL^-1^. Droplets (125 µL) of the suspension were dispensed into PLA lattice inserts placed in 12-well plates and allowed to gel at 37 °C for 30 min. Constructs were cultured in DMEM supplemented with 10% FBS, 1% non-essential amino acids, and 1% penicillin–streptomycin for 5 days, during which HepG2 cells formed compact spheroids within the hydrogel scaffold.

At day 0 of co-culture, devices were sterilized by UV irradiation for 60 min, and preconditioned Caco-2 monolayers (14 days), HSkM monolayers (2 days), and HepG2 spheroids (5 days) were integrated. Transwell inserts containing Caco-2 and HSkM were positioned in the designated side chambers, while HepG2 hydrogel constructs were placed in the central hepatic compartment. The assembled devices were mounted on a rocking platform (±6°, 10 cycles min^-1^) to generate gravity-driven bidirectional flow through the interconnecting circulation channel or maintained under static conditions. The circulation channel was filled with shared DMEM supplemented with 10% FBS, 1% non-essential amino acids, and 1% penicillin–streptomycin, while the apical compartments of Caco-2 and HSkM inserts were maintained with their respective culture media. All media were replenished every 24 h.

### Functional assays

Barrier integrity of Caco-2 monolayers was assessed by transepithelial electrical resistance (TEER) using an EVOM2 voltohmmeter (World Precision Instruments) with chopstick electrodes. Skeletal muscle metabolic activity was evaluated by measuring lactate concentration in the circulation channel using a colorimetric lactate assay (Dojindo) according to the manufacturer’s instructions.

For immunofluorescence (IF) assays, cells and spheroids were fixed with 4% paraformaldehyde for 20 min at room temperature, permeabilized with 0.1% Triton X-100 in PBS, and blocked with 5% bovine serum albumin (BSA) in PBS for 1 h. Samples were incubated overnight at 4 °C with the following primary antibodies: ZO-1 (rabbit anti-human; Proteintech) for tight junctions, albumin (ALB, mouse anti-human; Bioss Inc.) for hepatocytes, and myosin heavy chain (MYH, clone B-5, Alexa Fluor 594 conjugate; Thermo Fisher) for skeletal muscle. After washing, samples were incubated for 1 h at room temperature with species-appropriate secondary antibodies, including Alexa Fluor 488 goat anti-rabbit and Alexa Fluor 555 goat anti-mouse (Thermo Fisher). Nuclei were counterstained with DAPI (Thermo Fisher) before imaging. Imaging was performed using fluorescence microscopy (KEYENCE, Tokyo, Japan).

### Data analysis and statistics

For the investigation of cell morphology and quantification of corrected total fluorescence intensity (CTCF), images were analyzed using ImageJ software (National Institutes of Health, Bethesda, MD, USA) and CellProfiler software^25^ (Version 3.1.8; Broad Institute of Harvard and MIT, Cambridge, MA, USA). All quantitative data are presented as mean ± standard error of the mean (SEM). Statistical analyses were performed using one-way ANOVA with Dunnett’s multiple comparison test or unpaired two-tailed Student’s t-test, as appropriate. Graphs and statistical outputs were generated using GraphPad Prism 9 (GraphPad Software, San Diego, CA, USA).

## Acknowledgments

We acknowledge the Ritsumeikan Global Innovation Research Organization (R-GIRO). We thank Mr. Keita Watanabe, Department of Pharmaceutical Sciences, Ritsumeikan University for the assistance in culturing Caco-2 cells. This work was generously supported by the Japan Society for the Promotion of Science (24K15712), awarded to Rodi Kado Abdalkader.

## Authors contribution statement

Rodi Kado Abdalkader: Conceptualized and managed the project, designed, and performed experiments, analyzed, and interpreted data, visualized the data, and wrote the manuscript. Takuya Fujita: provided resources and contributed to manuscript refinement. All authors critically reviewed the manuscript and agreed with the publication.

## Additional information

Competing interests: All authors declare no competing financial interest.

