## Supplementary Information for "A 3D-printed pump-free multi-organ-on-a-chip platform for modeling the intestine–liver–muscle axis"

*Corresponding author.

[†]Earlier known as Rodi Abdalkader

**Fluid motion simulation**

Fluid motion was represented as a laminar, parabolic Poiseuille-like velocity profile with a time-dependent mean velocity defined as $U_{mean}(t)=U_{peak}\sin(2\pi ft)$, where $U_{peak}=5$ mm $s^{-1}$ was matched to the experimentally observed peak velocity, and f=10 cycles $min^{-1}$ represented the rocking frequency. The velocity profile across the channel height was assumed to be parabolic (like laminar Poiseuille flow), so the local velocity was calculated according to the equation:

$$\mathrm{u}(y,t) = \mathrm{U}_{\mathrm{mean}}(t) \cdot 1.5\left[1-\left(\frac{y-H/2}{H/2}\right)^2\right]$$

Where u(y,t) is velocity at vertical position y and time t, H is channel height (2 mm). The velocity is maximum at the center and zero at the walls.

Tracer transport was described by the two-dimensional advection–diffusion equation:

$$\frac{\partial \mathrm{C}}{\partial t} + \mathrm{u}(x,y,t)\frac{\partial \mathrm{C}}{\partial \mathrm{x}} = \mathrm{D}\left(\frac{\partial^2 \mathrm{C}}{\partial x^2} + \frac{\partial^2 \mathrm{C}}{\partial y^2}\right)$$

where C is the tracer concentration, $u(x,y,t)$ is the oscillatory velocity field, and D is the diffusion coefficient of fluorescein in water ($4\times10^{-10}$ $m^2s^{-1}$). No-flux boundary conditions were applied at the top and bottom walls, while approximate zero-gradient conditions were applied at the inlet and outlet.

The simulation domain was discretized into a 240 × 40 grid, and the time step was automatically adjusted to satisfy both Courant–Friedrichs–Lewy (CFL) stability and diffusion constraints. Initial tracer concentration was set as a uniform plug at the channel inlet. Numerical integration and visualization were implemented in Python using NumPy and Matplotlib, and the simulation output was rendered as an animation of tracer oscillation and mixing.

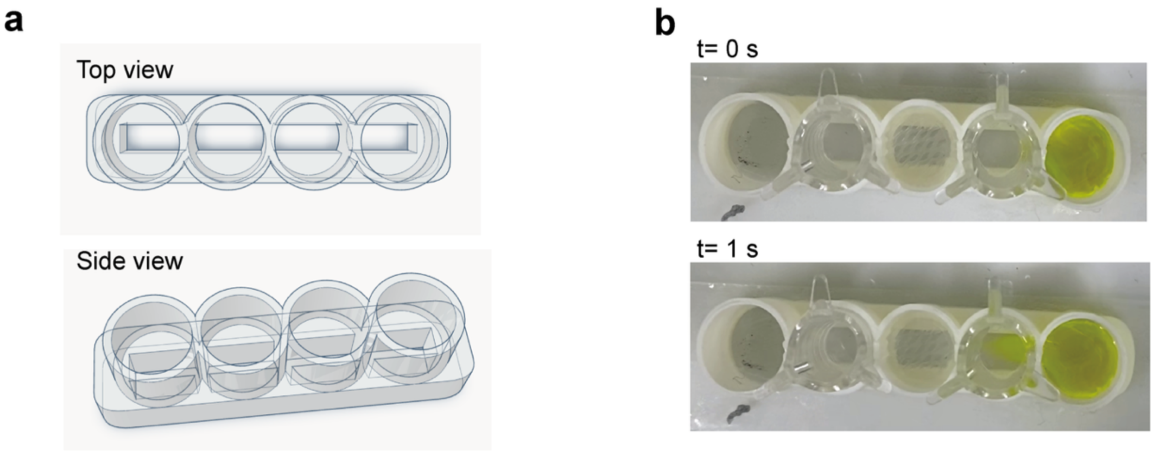


**Supplementary Figure 1. Design and flow characterization of the 3D-printed multi-organ-on-a-chip device.** **a**, Computer-aided design (CAD) schematics of the device showing the arrangement of five circular chambers and the connecting circulation channel (top and side views). The central chamber was designed to host hydrogel-embedded hepatic spheroids, while the side chambers accommodated Transwell cell culture inserts for intestinal and muscle compartments. **b**, Experimental validation of oscillatory flow using fluorescein sodium (1 mg $mL^{-1}$) under rocking conditions (±6°, 10 cycles $min^{-1}$). Representative images at t = 0 s and t = 1 s show dye displacement between adjacent chambers, confirming bidirectional perfusion without external pumps.

**Supplementary Code S1. Rocking-driven channel-flow and tracer transport simulation (Python)**

```
import numpy as np
import matplotlib.pyplot as plt
from matplotlib import animation
# Geometry & rocking params
L_mm, H_mm = 40, 2.0          # channel length & height in mm
L, H = L_mm/1000, H_mm/1000     # meters
f = 10.0/60.0                  # rocking frequency (Hz) = 10 cpm
theta0_deg = 6.0               # ±6°
# Target peak *mean* axial velocity (match experiment)
u_peak = 0.005                 # 5 mm/s (in m/s)
# Tracer diffusion (fluorescein ~4e-10 m^2/s)
D = 4.0e-10
# Simulation time
t_total = 20.0                 # seconds
# ------------------------
# Numerical grid
# ------------------------
Nx, Ny = 240, 40
x = np.linspace(0, L, Nx)
y = np.linspace(0, H, Ny)
dx = x[1] - x[0]
dy = y[1] - y[0]
# Time step from stability limits
cfl = 0.4
dt_adv = cfl * dx / max(u_peak, 1e-9)
dt_dif = 0.25 * min(dx, dy)**2 / D
dt = min(dt_adv, dt_dif)
nt = int(np.ceil(t_total / dt))
# ------------------------
# Initial tracer condition
# ------------------------
C = np.zeros((Ny, Nx), dtype=float)
C[:, :int(0.15*Nx)] = 1.0   # slug near inlet
# ------------------------
# Velocity profile: Poiseuille between plates
# u(y,t) = Umean(t) * 1.5 * (1 - ((y - H/2)/(H/2))^2)
# ------------------------
yy = y[None, :]
yc = H/2.0
parabolic = 1.5 * (1.0 - ((yy - yc)/(H/2.0))**2)
parabolic[parabolic < 0] = 0
def u_field(t):
    Umean = u_peak * np.sin(2*np.pi*f*t)
    return (Umean * parabolic.T) * np.ones((Ny, Nx))
# ------------------------
# One explicit advection–diffusion step with
# upwind in x and central diffusion; no-flux at y walls
# ------------------------
def step(C, t, dt):
    u = u_field(t)

    # Upwind in x
```

```
    Cx_fwd = np.roll(C, -1, axis=1) - C
    Cx_bwd = C - np.roll(C, 1, axis=1)
    adv_x = np.where(u >= 0, u * Cx_bwd/dx, u * Cx_fwd/dx)
    # Diffusion
    Cxx = (np.roll(C, -1, axis=1) - 2*C + np.roll(C, 1, axis=1)) / dx**2
    Cyy = (np.roll(C, -1, axis=0) - 2*C + np.roll(C, 1, axis=0)) / dy**2
    # No-flux y boundaries (Neumann)
    Cyy[0, :]  = (C[1, :]  - C[0, :])  / dy**2
    Cyy[-1, :] = (C[-2, :] - C[-1, :]) / dy**2
    # Gentle handling at x-ends
    adv_x[:, 0]  = adv_x[:, 1]
    adv_x[:, -1] = adv_x[:, -2]
    Cxx[:, 0]    = Cxx[:, 1]
    Cxx[:, -1]   = Cxx[:, -2]
    return C + dt * (-adv_x + D*(Cxx + Cyy))
# ------------------------
# Animation
# ------------------------
fig, ax = plt.subplots(figsize=(8, 2.4))
im = ax.imshow(C, extent=[0, L_mm, 0, H_mm], origin='lower', aspect='auto')
cb = plt.colorbar(im, ax=ax, label='Tracer (a.u.)')
ax.set_xlabel('x (mm)')
ax.set_ylabel('y (mm)')
ttl = ax.set_title('t = 0.00 s | Umean = 0.00 mm/s')
# Quiver (sparse)
qx_idx = np.linspace(0, Nx-1, 30, dtype=int)
qy_idx = np.linspace(0, Ny-1, 6, dtype=int)
Qx, Qy = np.meshgrid(x[qx_idx]*1000, y[qy_idx]*1000)
quiv = ax.quiver(Qx, Qy, np.zeros_like(Qx), np.zeros_like(Qy), scale=50, width=0.003)
frames = 400
substeps = max(1, nt // frames)
def animate(k):
    global C
    for _ in range(substeps):
        t = (k*substeps + _) * dt
        C = step(C, t, dt)
    im.set_data(C)
    u = u_field(k*substeps*dt)[qy_idx][:, qx_idx] * 1000.0  # mm/s
    quiv.set_UVC(u, np.zeros_like(u))
    Umean = u_peak * np.sin(2*np.pi*f*(k*substeps*dt)) * 1000.0
    ttl.set_text(f"t = {k*substeps*dt:5.2f} s | Umean = {Umean:5.2f} mm/s")
    return [im, quiv, ttl]
anim = animation.FuncAnimation(fig, animate, frames=frames, interval=40, blit=False)
# --- SAVE OPTION 1: MP4 (requires ffmpeg) ---
# anim.save('rocking_flow.mp4', dpi=150, writer=animation.FFMpegWriter(fps=25))
# --- SAVE OPTION 2: GIF (requires Pillow) ---
# anim.save('rocking_flow.gif', dpi=150, writer='pillow')
plt.tight_layout()
plt.show()
```
