## Supplementary figures and images for "A 3D-printed pump-free multi-organ-on-a-chip platform for modeling the intestine–liver–muscle axis"

### Supplementary video 1

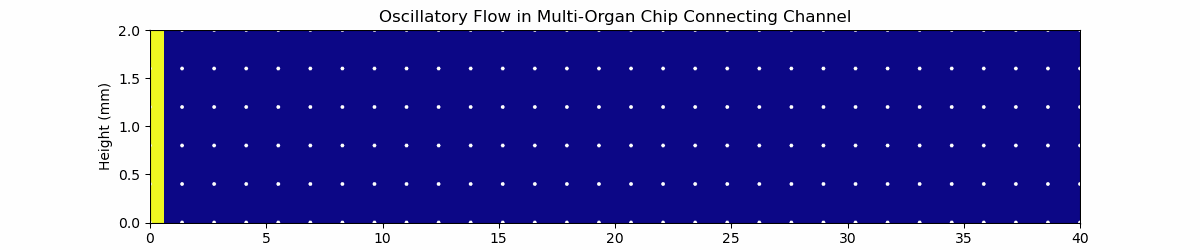
